# Clingen Cancer Somatic Working Group – standardizing and democratizing access to cancer molecular diagnostic data to drive translational research

**DOI:** 10.1101/212225

**Authors:** Subha Madhavan, Deborah Ritter, Christine Micheel, Shruti Rao, Angshumoy Roy, Dmitriy Sonkin, Matthew Mccoy, Malachi Griffith, Obi L Griffith, Peter Mcgarvey, Shashikant Kulkarni, On Behalf Of The Clingen Somatic Working Group

## Abstract

A growing number of academic and community clinics are conducting genomic testing to inform treatment decisions for cancer patients (1). In the last 3–5 years, there has been a rapid increase in clinical use of next generation sequencing (NGS) based cancer molecular diagnostic (MolDx) testing (2). The increasing availability and decreasing cost of tumor genomic profiling means that physicians can now make treatment decisions armed with patient-specific genetic information. Accumulating research in the cancer biology field indicates that there is significant potential to improve cancer patient outcomes by effectively leveraging this rich source of genomic data in treatment planning (3). To achieve truly personalized medicine in oncology, it is critical to catalog cancer sequence variants from MolDx testing for their clinical relevance along with treatment information and patient outcomes, and to do so in a way that supports large-scale data aggregation and new hypothesis generation. One critical challenge to encoding variant data is adopting a standard of annotation of those variants that are clinically actionable. Through the NIH-funded Clinical Genome Resource (ClinGen) (4), in collaboration with NLM’s ClinVar database and >50 academic and industry based cancer research organizations, we developed the Minimal Variant Level Data (MVLD) framework to standardize reporting and interpretation of drug associated alterations (5). We are currently involved in collaborative efforts to align the MVLD framework with parallel, complementary sequence variants interpretation clinical guidelines from the Association of Molecular Pathologists (AMP) for clinical labs (6). In order to truly democratize access to MolDx data for care and research needs, these standards must be harmonized to support sharing of clinical cancer variants. Here we describe the processes and methods developed within the ClinGen’s Somatic WG in collaboration with over 60 cancer care and research organizations as well as CLIA-certified, CAP-accredited clinical testing labs to develop standards for cancer variant interpretation and sharing.

## ClinGen

To address these needs of capturing, standardizing and sharing clinically relevant variants, the Clinical Genome Resource, ClinGen (4) collaboration was established in 2012 and has been developing interconnected community resources to improve our understanding of genomic variation and enhance its use in clinical care. ClinGen represents a strong partnership among public, academic, and private institutions that relies on collaboration between the NIH, academic and commercial laboratories operating in both the research and clinical realms. ClinGen is also engaging numerous entities, including professional societies, to ensure that the resources that are produced meet community expectations. The Somatic working group (Somatic WG) is a clinical domain working group within the ClinGen consortium and was established in 2015 to address standardization and sharing of cancer MolDx test results described here.

## The Standard

In order to standardize the collection of clinically relevant somatic data, the Somatic WG of ClinGen created a framework of consensus data elements titled "Minimum Variant Level Data” (MVLD) (5). MVLD was developed with input from multiple stakeholders ranging from database engineers to researchers and somatic clinical laboratory directors, as well as input from multiple current databases that collect cancer variant data. Briefly, MVLD consists of three sections: allele descriptive, allele interpretive and somatic interpretive. The allele descriptive section contains data elements that describe the genome position, gene, chromosome, genomic location, reference transcript and protein. The allele interpretive section contains data elements describing the somatic classification (confirmed somatic, confirmed germline or unknown), the DNA and protein substitution, the variant type and consequence and PubMed identifiers associated with interpretation. The somatic interpretive section contains the most clinically relevant data, and is the section that required the most discussion and consensus-building among the working group members. The somatic interpretive section contains a description of the cancer type (NCI Thesaurus, Oncotree, Disease Ontology), the Biomarker Class (Diagnostic, Prognostic, Predictive), the Therapeutic Context (associated drugs), Effect (Resistant, Responsive, Not-Responsive, Sensitive, Reduced-Sensitivity), Level of Evidence (a tiered system similar to the recent AMP/CAP/ASCO guidelines) (6) and Sub-Level of Evidence (reporting of trials, metadata analysis, preclinical data or inferential data). Readers are referred to the publication for a more detailed description of these data elements.

Since the publication of MVLD, recent guidelines on somatic variant interpretation have been published through a joint effort of the Association for Molecular Pathology (AMP), College of American Pathologists (CAP) and the American Society of Clinical Oncology (ASCO)(6). We intend to fully harmonize MVLD elements with these guidelines; mapping any specific criteria to the current version of MVLD and revising MVLD to accommodate new elements. There are distinct areas of agreement between MVLD and AMP/CAP/ASCO guidelines, such as using HUGO-approved nomenclature and HGVS formatting for variants. However, there are also sizable and nuanced differences that need resolution to sync the guidelines with the MVLD data structure.

One area of immediate critical harmonization needed is in the Somatic Interpretive Level of Evidence and Sub-Level of Evidence in MVLD, which was drawn from the Cancer Driver Log (CanDL) (7). The AMP/CAP/ASCO guidelines contain classification for uncertain (Tier III) and benign (Tier IV) variants, while MVLD was not initially designed to incorporate these types of variants. However, the necessity and relevance of uncertain or benign variants is apparent in that they too can aid clinical diagnosis. The AMP/CAP/ASCO guidelines Tier I Level A and MVLD Tier 1 are the same, but AMP/CAP/ASCO further provides Level B to sustain interpretations that derive from well-established studies that are not yet FDA or NCCN approved. Similarly, there are numerous nuanced differences between AMP/CAP/ASCO Tier II Level C and D and MVLD Tier 2, 3 and 4. The Sub-Level of Evidence in MVLD is incorporated in AMP/CAP/ASCO at various Tiers as well. Instead of partially modifying the MVLD Level of Evidence and Sub-Level of Evidence, we propose to absorb the Sub-Level of Evidence element into the Level of Evidence and to fully adopt the classification system proposed by AMP/CAP/ASCO into the Somatic Interpretive Level of Evidence shown in **Figure 1**. Cancer-type might be agnostic, for example with PD-L1 testing for Keytruda.

**Fig.1.**
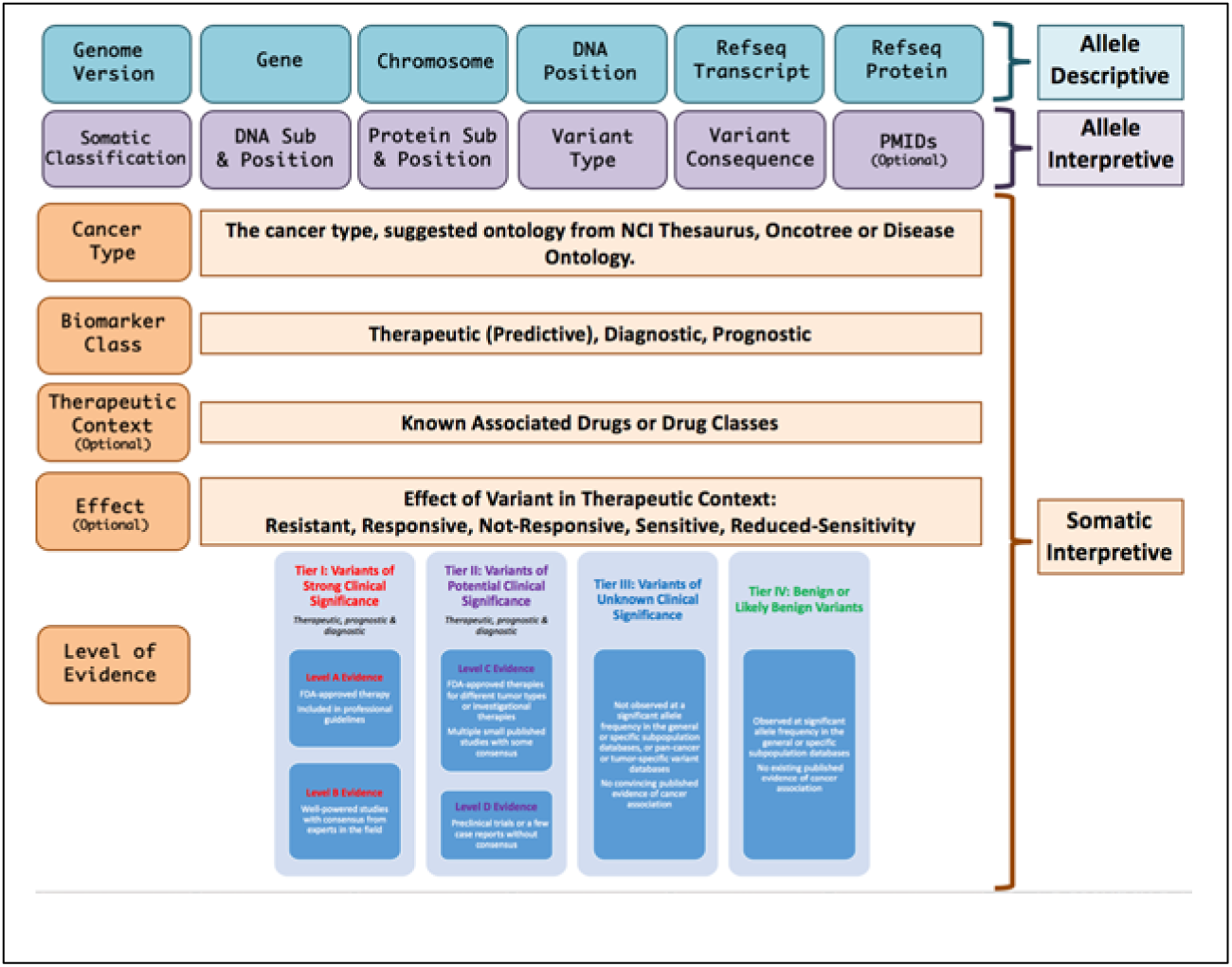
Mapping AMP guidelines to the ClinGen MVLD framework

In addition, there are other moderate differences. AMP/CAP/ASCO suggested the variant allele fraction of the somatic variant was important to report, and MVLD did not include a field for this. In general, MVLD did not focus on negative results or include fields for lack of supporting variant evidence, but will consider doing so in future revisions by incorporating them into the Tier III and Tier IV categories. For Biomarker Classifications, AMP/CAP/ASCO uses the term "therapeutic" which is similar to the MVLD "predictive"; we are proposing to adopt "therapeutic" as well, and will start by using a combination of both "therapeutic/predictive" to ensure clarity. AMP/CAP/ASCO did not touch upon standardized ontology for cancer type, while MVLD proposed use of NCI Thesaurus, Oncotree or SNOMED. We will be revising MVLD to adopt Disease Ontology as well due to its popularity and prevalence.

The Somatic WG of ClinGen plans to continue reviewing the above detailed distinctions and again use a consensus-driven, round table approach to resolve and revise the MVLD to accommodate and support somatic variant interpretation guidelines. While the AMP/CAP/ASCO somatic interpretation guidelines have been very well received, consistent feedback has promoted discussions of increased granularity for the guidelines, similar to the germline variant interpretation guidelines of the American College of Medical Genetics and Genomics (ACMG). The Somatic WG of ClinGen is preparing a joint effort with ACMG and AMP/CAP/ASCO to incorporate this level of detail into somatic interpretation guidelines.

## Variant Curation SOP and expert review

Our variant curation and interpretation process leverages the strengths of ClinGen Somatic WG, the consortium of multi-disciplinary experts in somatic variants in cancers, CIViC (8), a cancer variant knowledgebase and crowdsourced curation system and ClinVar (9), an NCBI submission-driven database for variants. ClinGen brings/develops organized clinical and biomedical expertise, best practices and SOPs, CIViC provides a curation interface and interpretation portal, and ClinVar allows widespread dissemination of the expert-curated content and provides patient-level observations of variants in clinical settings back into ClinGen/CIViC.

The ClinGen Somatic variant curation and expert review process is shown in Figure 2. New submissions or revisions are made through data entry pages in CIViC that support dynamic form adjustments, live type-ahead suggestions, ontology look-ups, and warnings for merge conflicts. Discussion pages track the complete history of comments and revisions. Curators and editors have the option to “follow” any entry (gene, variant, evidence) to receive notifications of comments, proposed changes or additions. Curators can also communicate with others in the CIViC community directly through site mentions, updates and messages. All curated entries can be “flagged” for problems or revisions can be proposed. Flagging allows for easy marking of content needing immediate review or users can flag entries which require more caution with use as diagnostic markers, while revisions are tracked and displayed with detailed GitHub-style diffs and comments.

**Fig. 2.**
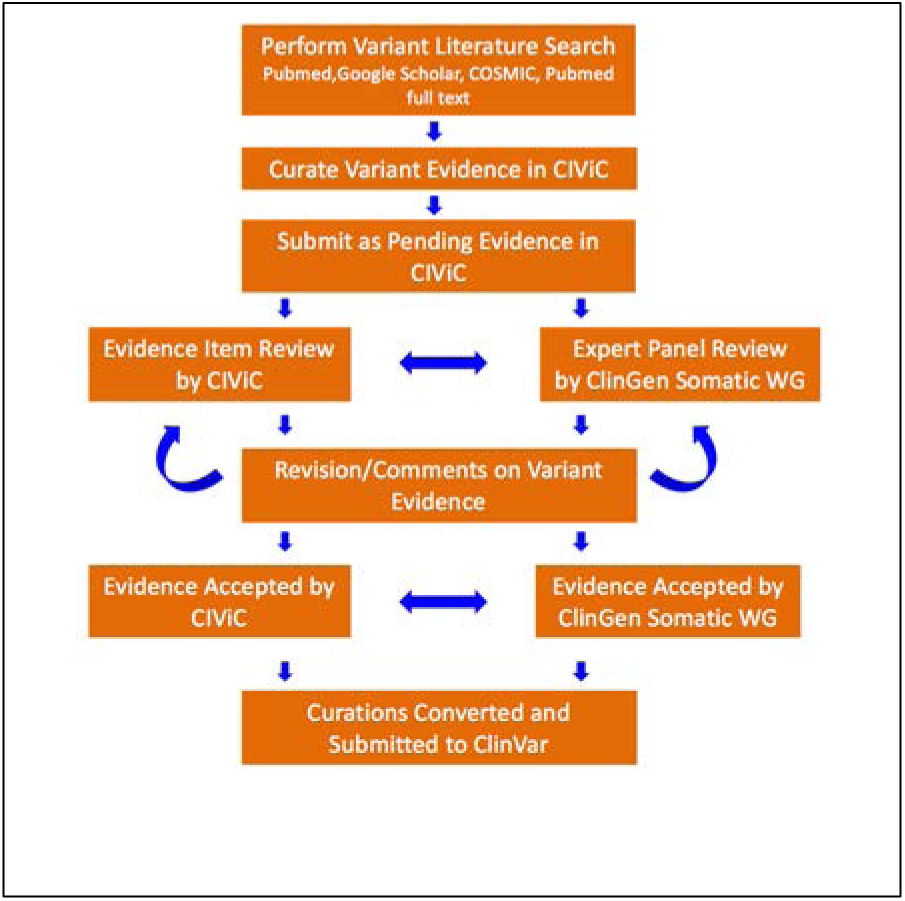
ClinGen Somatic Variant Curation SOP

Curators can create detailed profiles so that their efforts are recognized by awarding badges for curation activity milestones to encourage and recognize participation. Curators can also join formal curation organizations within the CIViC community, for example the ClinGen Somatic Working Group exists as a CIViC organization, currently with 12 active members.

The high quality of CIViC content is encouraged through several mechanisms. First, content creation is completely transparent with detailed provenance for all additions and revisions. Problems can be identified quickly by publicly accessible comment or flag features. Anyone can become a curator with the ability to flag, comment, or submit new/revised content. To ensure all content is reviewed by at least 2 CIViC reviewers, new entries must be reviewed by a site editor and users cannot accept their own contributions. There is an additional layer of expert review by the ClinGen Somatic WG disease or gene centric taskforces, which are described in Community Engagement.

## MolDx2MVLD mapping tools

To complement the crowdsourced expert variant curation process, members of the ClinGen WG are designing and implementing tools to support mapping of clinical MolDx data to standards and automated importation of this data into research databases, for example, G-DOC (10) (Georgetown Database of Cancer) and SEER (11) to drive new hypothesis generation for translational research. The primary goal of this tool is to enable broad sharing of de-identified MolDx data from clinical laboratories for novel hypothesis generation and evidence collection for clinical actionability.

The general components of the tool to map MolDx data to the MVLD system are outlined in Figure 3. The four main components of the tool are: 1) ETL (Extract, Transform, Load) tools to parse individual sources, extract and format information required for MVLD descriptive and interpretive elements; 2) MVLD Mapper to map the extracted information to the MVLD standard, harmonize the elements to standard identifiers and ontologies used in public data repositories, identify missing data elements, and attempt to fill in missing values if possible; 3) a simple QA/QC interface for checking results and correcting or adding missing values; and 4) MVLD formatter to output the information in various formats, e.g., xml, tabular, or others to interface with EHRs and translational research databases depending on user needs.

**Fig. 3.**
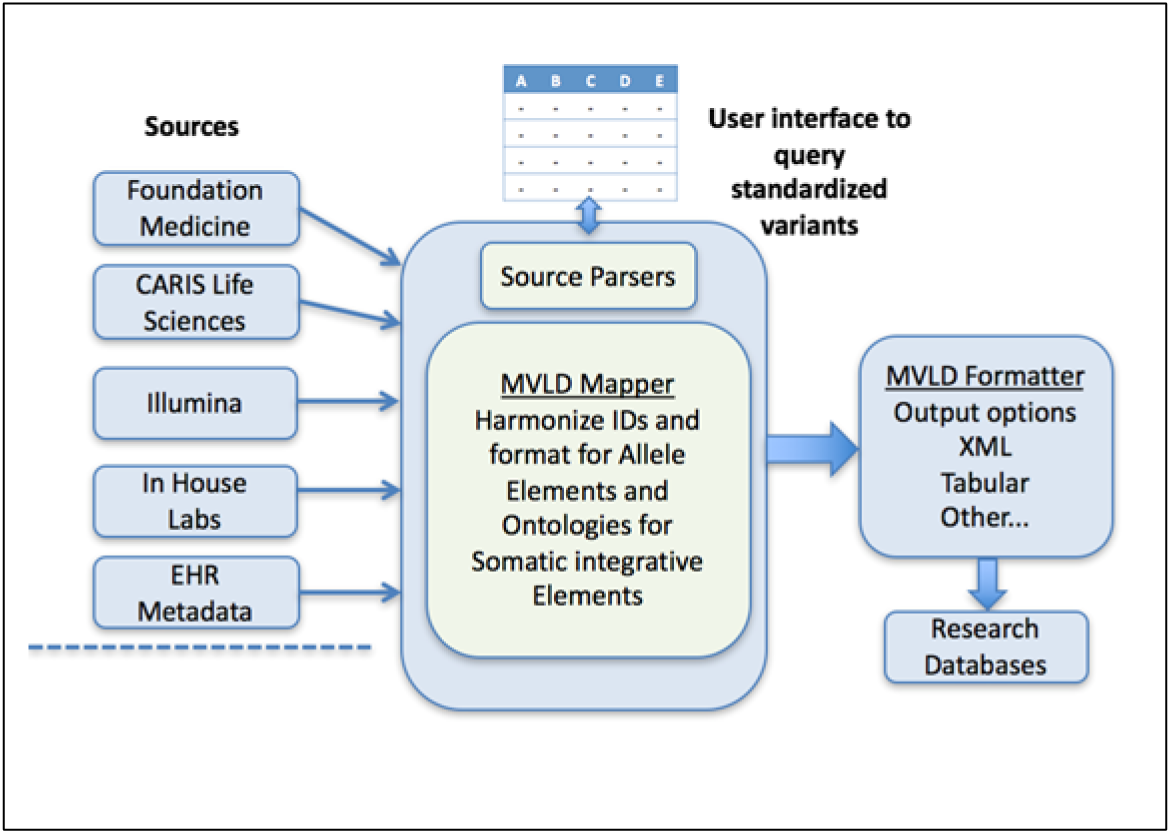
Cancer MolDx – MVLD software schematic

### ETL tools

MolDx laboratory sources use different internal formats and standards to identify a variant and describe the results of their tests. For example, some labs use only gene names and protein changes as a description; others provide transcript identifiers and the specific DNA change in the transcript and/or the exact chromosomal location. To date we have not seen a report using a complete HGVS (Human Genome Variation Society) formatted sequenceID+variation name, though some labs provide the information to create the description. There are also differences in how data is distributed, with some labs providing results in tabular formats while others provide XML. We will create parsers and logic for each source to extract and perform initial transformations. **Figure 4** shows an example of an XML excerpt from the Foundation Medicine report on a patient with breast carcinoma. This patient’s molecular testing identified a variant E545K in the PIK3CA gene with potential benefit from mTOR inhibitors such as Everolimus or Temsirolimus. The figure shows mapping of XML output from the lab to elements in the MVLD standard. Similar mapping is being conducted for all 18 data elements in MVLD to various commercial labs. Lab formats to integrate are prioritized by our stakeholder community based on most widely used labs.

**Fig. 4.**
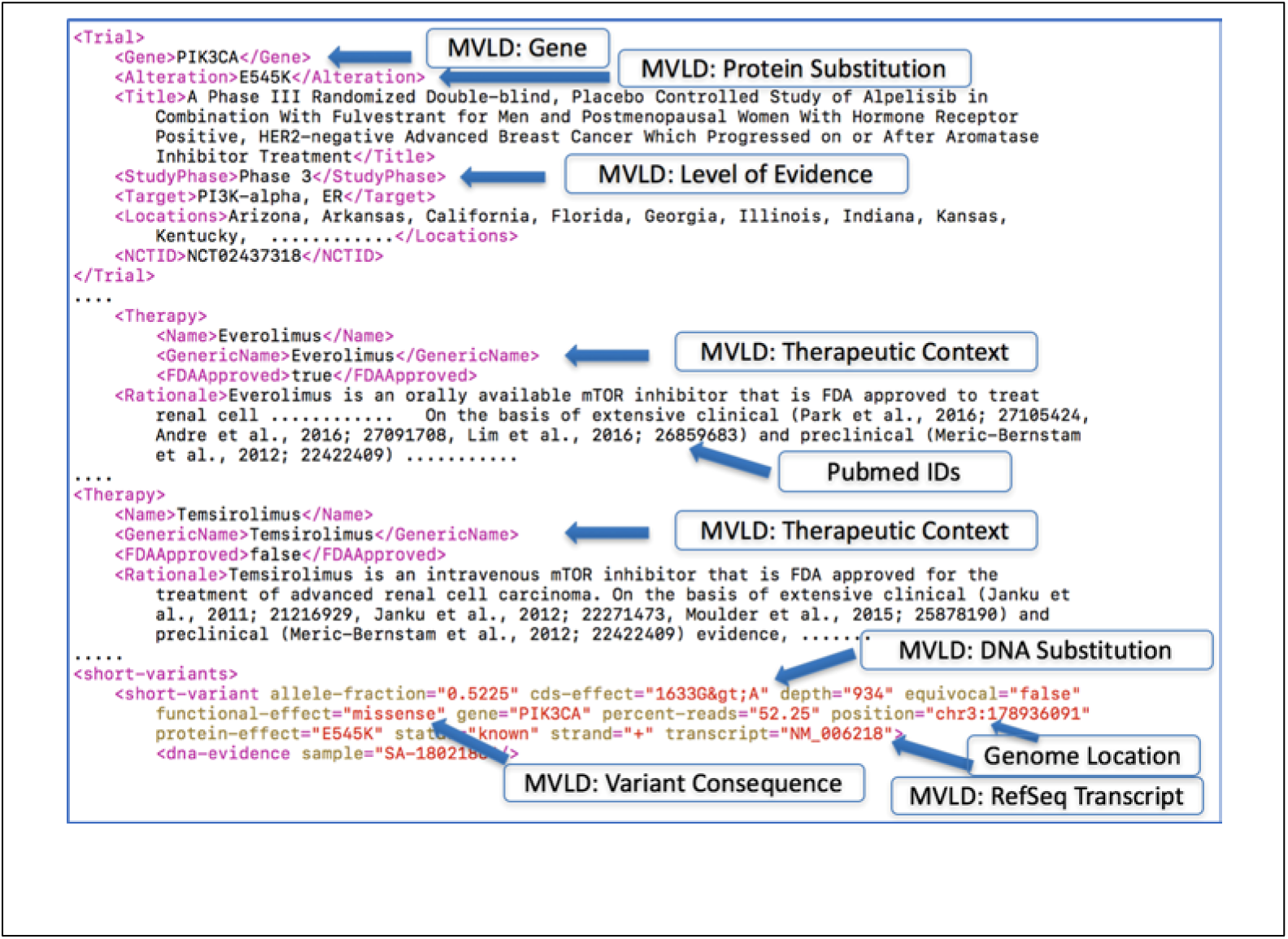
A sample mapping of FM XML file to MVLD standard is shown. Not all lab reports are as well structured

### MVLD mapper

This is the core component of the system. We expect variations in how the descriptive and interpretive properties in MVLD will be expressed, and the mapper attempts to harmonize these representations to the most informative common representation using variant-specific APIs connected to international variation databases and ontology servers. For allele properties, we use NCBI’s E-utilities including their new variation API services that allow users to compare and return all equivalent alleles using multiple NCBI identifiers including a canonical identifier (12). The tool first checks the ClinVar database to see if a representation of the variation exists or the test is registered with the Genetic Testing Registry, in which case ClinVar may contain all the variants tested (13). ClinGen has released an Allele Registry with APIs that also attempts to link equivalent variant alleles to a canonical representation but is not NCBI centric in that ENSEMBL IDs and even ExAC alleles are supported and novel alleles can be submitted (14).

MVLD Interpretive elements like cancer type can be standardized using APIs for terminology servers. The WG identifies terminology standards used in key data fields in lab reports, such as disease and drug names, ICD codes, or the 10-digit national drug code and map them to MVLD-recommended standards. If any term is unknown, the MolDx processor attempts to automatically map to terminologies in the NCI Thesaurus (NCIt) using LexEVS Terminology Server APIs or BioPortal APIs (15, 16). In all cases, the original value is stored along with the selected mappings to defined terminologies. This allows the data to be in a uniform format that follows standards and ontologies, while allowing for more integrated search functions within systems like ClinVar.

### MVLD Formatter

The formatter module provides multiple output options for target research databases. The Initial output will be delimited tables or XML mainly for consumption by institutional databases (EHRs) that want to store the MVLD standardized data. We will work with the community to define and build in additional XML or other formats (e.g. JSON) from labs.

## Community Engagement

Many standards and frameworks fail due to lack of community engagement and adoption. To avoid this, the ClinGen clinical domain working group engaged both strategic leaders and tactical implementers from over 60 cancer centers, industry partners and federal agencies. These include active participation from organizations such as Georgetown University Lombardi Cancer Center, Baylor College of Medicine, Vanderbilt University Medical Center, Washington University School of Medicine, Moffitt Cancer Center, Illumina, Molecular Match, NCI, NHGRI and FDA. An initial survey of participating organizations identified major challenges in somatic variant assessment, clinical interpretation pipelines and open tools for variant curation and expert review. The survey results also indicated the use of a tiered system of variants for clinical actionability (FDA-approved/NCC guidelines, clinical trials data, pre-clinical data, mechanistic/pathway level evidence).

These results motivated the efforts to develop the MVLD to help standardize how clinical labs report MolDx data to patients, clinicians and regulatory agencies. We also engaged members from AMP (Association of Molecular Pathologists) and CAP (College of American Pathologists) somatic practice guideline committees to help drive adoption of MVLD within their professional societies and members. ClinGen Somatic WG is also actively working with Global Alliance for Genomic Health (GA4GH)’s Variant Interpretation for Cancer Consortium (VICC). The VICC seeks to integrate global efforts for the clinical interpretation of cancer variants. The ClinGen Somatic WG engages various experts in the cancer research and care communities through taskforces. These taskforces are self-organized expert groups in a particular cancer type, gene or a pathway. Three such taskforces that are currently operational are described below with other taskforces being routinely formed.

### Pediatric somatic taskforce

Cancers in children differ from those in adult patients in at least three key aspects. First, a number of childhood cancers occur exclusively in children and adolescents. The genomic landscape of such tumors, including medulloblastoma, neuroblastoma, and Wilms tumors, have been the focus of several recent large-scale research projects, such as the Therapeutically Applicable Research to Generate Effective Treatments (TARGET) (17), the Pediatric Cancer Genome Project (PCGP) (18), the Peds-MiOncoSeq (19) and the BASIC3 (20) projects, and have been found to harbor not only shared but several novel recurrent alterations when compared to the most common genomic alterations seen in adult epithelial malignancies. Such alterations include, in addition to SNVs and indels, a higher preponderance of gene fusions and copy number alterations. Second, genomic profiling of certain childhood cancers with particularly poor outcomes and with well-recognized adult counterparts such as glioblastoma have revealed remarkable non-overlap in the key genetic drivers and signaling pathways between the same disease histology in the two age groups (21). These studies highlight the diversity of genomic alterations that are exclusive or enriched in childhood cancers. Assessment of the extent of curation of key childhood cancer genes and variants in CIViC reveal many of the most common childhood cancer specific genes (e.g., *H3F3A*, *HIST1H3B*, *ACVR1*, *ATRX*, *DDX3X, DROSHA, DICER*) and variants to be absent or sparsely curated in CIViC and other cancer knowledgebases highlighting the urgent need for expert curation of novel childhood cancer genes.

Third, even when childhood tumors are found to harbor therapeutically targetable alterations that have been validated in clinical trials of adult patients, the feasibility and efficacy of such treatment options in children are often unclear; expert curation of such ‘actionable’ alterations in childhood cancers will require the insight of pediatric oncologists and other domain-specific experts in pediatric oncology.

The ClinGen Somatic Pediatric taskforce is addressing these issues through the expert variant curation and adjudication process described above.

### Pancreatic somatic taskforce

Pancreatic ductal adenocarcinoma (PDAC) is one of the most difficult cancers to treat, as physiological hurdles to preventative monitoring and an absence of early detection biomarkers result in many late stage diagnoses. The more advanced disease is typically metastatic and chemoresistant, with a 5-year survival is around 8% (22). The lack of noninvasive, PDAC specific biomarkers highlights the need to identify novel molecular signals associated with onset and progression, and to develop a better understanding of the role of somatic variation to improve individualized therapy decisions. The last stage diagnosis of PDAC typically results in the convergence of mutations in several known oncogenes and tumor suppressor; mutations activating KRAS are found in 95% of cases, inactivation of TP53 occurs in 75% of cases, and CDKN2A is inactivated in 95% of cases (23). These genes are known for their regulatory influence on cell proliferation and are particularly relevant to regulating a shift in active metabolic pathways promoting growth in a hypoxic microenvironment. This metabolic reprogramming found in PDAC not only increases glucose uptake and enhances glycolysis (24), but also increase the expression of glutamine metabolic pathways and promote production of NADPH/NADP+ (25). These cellular mechanisms are adapted to meet the energy requirement for cell division at a reduced oxygen consumption rate and represent an emerging set of PDAC targets that require somatic curation (26).

The occurrence of somatic variants in PDAC which overlap with known oncogenes and tumor suppressors provide the opportunity for targeted therapies which have proven effective in other cancers. In particular, a mutator phenotype of PDAC associated with a high mutation burden is driven by mutation to mismatch repair genes (27), and may benefit from any number of therapies developed to target cancers with DNA repair deficiencies. By extension, somatic variants found in other PDAC subtypes could benefit from successful advances in other cancerous tissues, and underscores the need for their high quality curation. Somatic variants in PDAC are being cataloged as part of the PANCAN Know Your Tumor program(1), and to date have identified 5473 unique variants from 432 genes.

Overall, the variants remain uncurated with about 38% of genes represented in CIViC, covering only 1–2% of variants observed in PDAC. Clearly there is a significant need for the expert curation of novel PDAC somatic variants, and the potential for improvement over current therapy options.

### TP53 taskforce

Tumor suppressor genes play important roles in cancer biology and could be inactivated by number of different mechanisms, such as deletions, insertions, inversions, loss of function point mutations, and loss of expression. Tumor suppressor gene TP53 is one of the most frequently altered genes across multiple tumor types. To fulfil its proper biological function four identical TP53 polypeptides must form a tetramer which functions as a transcription factor. Mostly due to this underlying molecular biology majority of TP53 inactivating alterations are loss of function point mutations. Majority of the loss of function point mutations are concentrated in the DNA binding domain and have different degree of dominant negative phenotype. MDM2-TP53 interaction inhibitors efficacy is currently under investigation in multiple clinical trials. Inactivating TP53 mutations prevent on target activity and efficacy of MDM2-TP53 interaction inhibitors, therefore only patients with intact TP53 could benefit from such inhibitors. Using preclinical (28) and clinical publications (29) on MDM2-TP53 interaction inhibitors efficacy in relation to TP53 status corresponding evidence items have been submitted to CIViC. At this point in time CIViC does not support entry of information on pathogenicity of somatic alterations; however this functionality will be added in the future. Due to current lack of ability to enter pathogenicity information MDM2-TP53 interaction inhibitors related evidence items for individual alteration have been limited to the ones which are directly mentioned in Saiki et al. 2015 publication (30).

## Future directions

A future goal of the ClinGen Somatic Working Group is to review and harmonize existing guidelines and guideline efforts related to curation, interpretation, and reporting of somatic alterations in cancer to provide a unified guideline to clinical labs to represent and interpret cancer variants. The working group is forming a task force that will bring together representatives from ClinGen, ACMG, AMP, ASCO and CAP, among other relevant organizations. This harmonization task force will seek to build upon recently published work such as the AMP/ASCO/CAP guidelines and the ClinGen Somatic Working Group’s MVLD for curation of somatic alterations in cancer. Further, this task force will work to make its standard compatible with other related guidelines, such as the ACMG/AMP guideline for interpretation of germline variants (31), and an ongoing effort within ACMG on the interpretation of copy number variants in neoplastic diseases. The task force will review and include work such as the Variant Interpretation in Cancer Consortium and AACR GENIE’s efforts to describe and standardize curation practices in the somatic cancer space (32).

Finally, the task force will review and include efforts of somatic cancer knowledgebases to map terminologies and levels of evidence schemes across knowledgebases (e.g., CIViC, OncoKB, PMKB, JAX-CKB, CGI, PCT, CanDL, G-DOC). The ClinGen gene and disease focused taskforces will provide usecases and variant examples to this harmonization effort, enabling a continually updated, unified guideline for somatic variant interpretation agreed upon by experts and serving the cancer MolDx community in their mission to democratize access to these important clinical datasets.

## Acknowledgements

Authors acknowledge support from the ClinGen grant NIH U01 HG007437. CIViC is supported by NIH U01 CA209936. MG is supported by NIH R00 HG007940. OLG is supported by NIH K22 CA188163.

